# Repertoire-Based Diagnostics Using Statistical Biophysics

**DOI:** 10.1101/519108

**Authors:** Rohit Arora, Joseph Kaplinsky, Anthony Li, Ramy Arnaout

## Abstract

A fundamental challenge in immunology is diagnostic classification based on repertoire sequence. We used the principle of maximum entropy (MaxEnt) to build compact representations of antibody (IgH) and T-cell receptor (TCRβ) CDR3 repertoires based on the statistical biophysical patterns latent in the frequency and ordering of repertoires’ constituent amino acids. This approach results in substantial advantages in quality, dimensionality, and training speed compared to MaxEnt models based solely on the standard 20-letter amino-acid alphabet. Descriptor-based models learn patterns that pure amino-acid-based models cannot. We demonstrate the utility of descriptor models by successfully classifying influenza vaccination status (AUC=0.97, p=4×10^-3^), requiring only 31 samples from 14 individuals. Descriptor-based MaxEnt modeling is a powerful new method for dissecting, encoding, and classifying complex repertoires.

## Introduction

A major challenge in systems immunology is determining how to describe the sequence-level heterogeneity of antibody (immunoglobulin; Ig) and T-cell receptor (TCR) repertoires in ways that facilitate the identification of meaningful patterns. Sequence-frequency distributions—for example, counts of unique IgH or TCRβ CDR3s—are commonly used but not ideal for interpersonal comparisons, since repertoires from different people are largely disjoint (Robins et al., 2010; Arnaout et al., 2011). Motif-frequency distributions, which count how often each of the 20” possible n-mers appears in a repertoire (for some choice of n), are more likely to overlap between individuals, but may fail to detect probabilistic or higher-order patterns and are subject to sampling-related bias unless *n* is small. Comparisons of frequency distributions between repertoires from different individuals have yielded important insights (Parameswaran et al., 2013; Kaplinsky et al., 2014; Emerson et al., 2017; Sun et al., 2017) but the limitations of this approach suggest a need for complementary methods. One such method is maximum-entropy (MaxEnt) modeling (Fig. 1).

**Figure 1.**
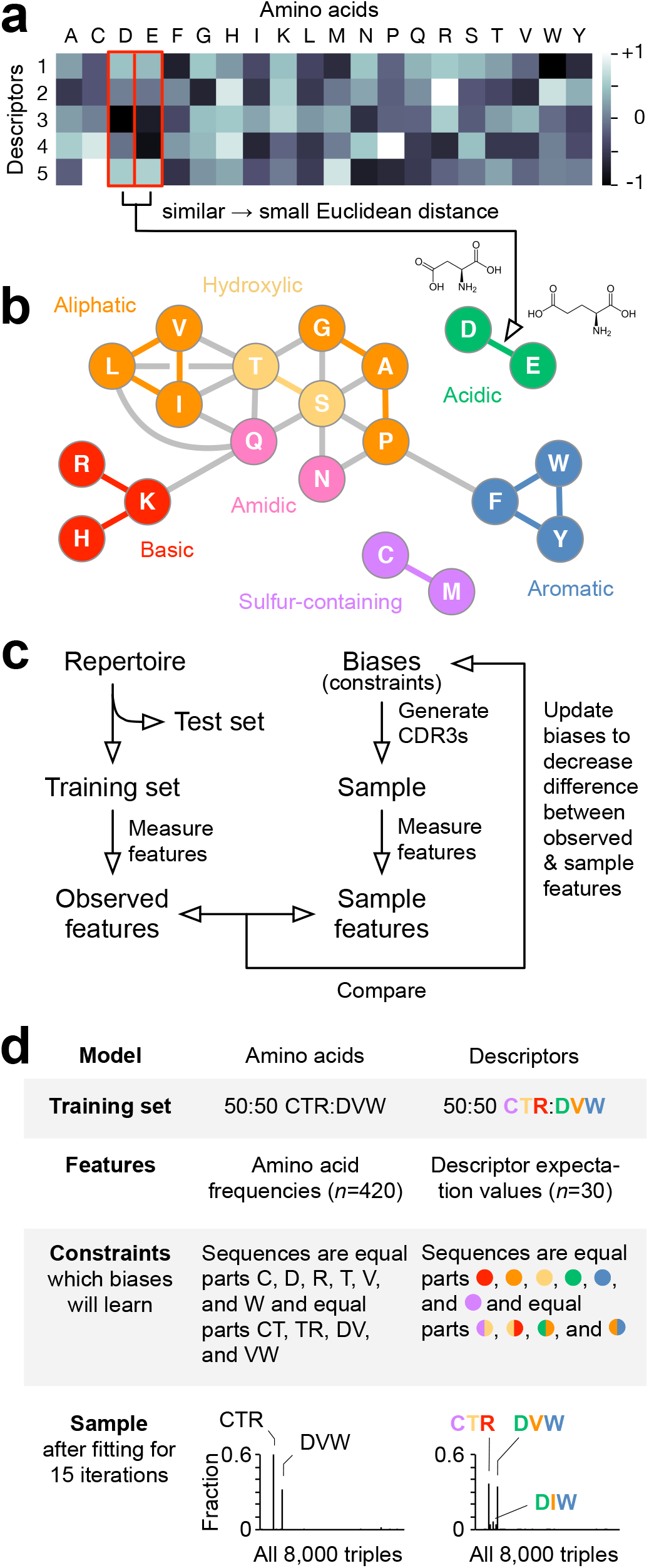
MaxEnt Based on Amino Acids’ Biophysical Properties. **(a)** Amino acids as vectors, shown here as a heatmap, in a 5-dimensional descriptor space. **(b)** Amino acids with similar properties lie near to each other in descriptor space. These similarities can be visualized by calculating all pairwise Euclidean distances of the amino acids in descriptor space, constructing a (complete, *K*_20_) network with the amino acids as nodes and the distances as weighted edges, and then for clarity keeping only edges with weights ≤1.1. For example, aspartate (D) and glutamate (E) (red boxes in (a)) lie near to each other in descriptor space, illustrated by their similar pattern in the heatmap (with prominent differences only in the dimension corresponding to descriptor 4), and so are adjacent in the network. Amino acids are colored according to a familiar groupings (basic, aliphatic, etc.) to demonstrate that their configuration in descriptor space agrees with these groupings. (c) Data preparation and model training. Repertoires were first split into training and test sets, and the features of the training set measured. Models were trained through iterative sampling, comparing of sample and observed features, and updating biases. **(d)** Example using a highly simplified toy repertoire consisting of a training set of two unique 3-amino-acid sequences, CTR and DVW (common in stems of IgH CDR3s). The models learn constraints that distinguish the training set from random 3-mers. For amino-acid models, constraints are the frequencies of letters; for descriptor models, constraints are expectation values of descriptors and descriptor products at given positions (here, nearest-neighbor pairs). Model output is shown in the last row. The descriptor model has learned the pattern of biophysical relationships, such that sequences that are biophysically similar to sequences in the training set also appear in the sample, albeit at lower frequency than the sequences in the training set.

MaxEnt models, which were first developed for statistical physics and information theory (Jaynes, 1957), can be used to describe repertoires (or other complex ensembles of proteins, nucleic acids, etc.) in terms of constraints called *biases* that determine the ways in which a given repertoire differs from a uniform distribution of sequences (Yeo and Burge, 2004; Russ et al., 2005; Seno et al., 2008; Mora et al., 2010; Marks et al., 2011). Given a set of *features*—for example, the frequencies of the 20 amino acids and the 20^2^=400 nearest-neighbor amino-acid pairs (“neighbors” being defined as contiguous N-to-C-terminus amino acids)—a MaxEnt model describes the degree to which each feature is biased away from its value in a uniform repertoire, taking all the other biases into account. For example, the bias for the pair cysteine-alanine (CA) describes the extent to which the frequency of CA in the repertoire differs from what would be expected given the frequencies of the individual amino acids C and A, the pairs XC and AX (for all amino acids X), and so on. MaxEnt models deconvolute the hundreds or thousands of interactions among features into separate components, which then together govern the generation of the observed sequence- and motif-frequency distributions. Thus MaxEnt models can be thought of as capturing the underlying generative structure of the repertoire.!

MaxEnt modeling of IgH and TCRβ CDR3s, as well as of other protein families, has shown that the frequencies of a single set of neighboring amino-acid pairs capture a remarkable amount of information (Russ et al., 2005; Seno et al., 2008; Mora et al., 2010; Marks et al., 2011), but not all of it (Bialek and Ranganathan, 2007). Additional sets of pairs—for example, second-, third-, or fourth-nearest neighbors (Mora et al., 2010)—add precision but at the cost of a substantial increase in the number of model parameters (400 per set of pairs). This increase can affect the coverage per feature (the total number of instances of the features in the sample divided by the number of features), model quality and interpretability, and training time. The root of the problem is that the amino-acid alphabet has 20 letters: as a result, parameters, data, and computational requirements scale roughly as powers of 20. The alphabet also causes a second important problem: letters in and of themselves, while a familiar and useful shorthand, lack information about similarities and differences among the multi-faceted biophysical entities they represent— the amino acids—for example, that A is more like glycine (G) than tyrosine (Y)—that may well contain meaningful patterns that are not obvious from, or captured by, the shorthand alone.

These two problems can be addressed simultaneously by swapping the traditional amino-acid alphabet for a smaller set of *descriptors* derived from amino acids’ physicochemical properties, especially for pairs and higher-order associations (e.g. consecutive triples) (Fig. 1a). Over two dozen lipophilic (e.g., hydrophobicity), steric (e.g. molecular weight) and electrical (e.g. charge) properties have been precisely measured (Sandberg et al., 1998; Kim et al., 2016). These properties have been shown to correlate with each other, such that the first few principal components (PCs) explain a majority of the overall variance (Hellberg et al., 1987; Sandberg et al., 1998). These PCs are natural candidates for a reduced alphabet: they define orthogonal dimensions of a continuous space in which the discrete amino acids are embedded (Fig. 1a). Whereas in “letter space” there is no concept of distance between amino acids, in “descriptor space” amino acids with similar properties are closer together (e.g., with A nearer G than Y) (Fig. 1 b). Such embeddings have been explored in immune-repertoire analysis (Greiff et al., 2017; Ostmeyer et al., 2017, 2019) and other contexts (Dosztányi and Torda, 2001; Walter et al., 2005; Susko and Roger, 2007; Stephenson and Freeland, 2013). We investigated whether descriptor-based MaxEnt models of IgH and TCRβ CDR3 repertoires could improve on models based on amino acids alone by allowing more data per parameter (less sampling error), shorter training time, and better interpretability (Fig. 1c-d), in principle leading to better models useful for classification of states of health and disease.

## Results

Using 26 measurements carried out on the 20 standard amino acids, we derived five biophysical descriptors that together explained 92% of the variance in amino acids’ physicochemical properties. Each descriptor is a PC, i.e. a linear combination of the measurements. The first three descriptors corresponded roughly to surface area/chromatographic properties (explaining 41% of the overall variance), van der Waals volume (25%), and charge (14%) and together explained 79% of variance, an increase over the 68% previously reported for the first three descriptors derived from measures of both the standard and additional non-canonical amino acids (Sandberg et al., 1998).

We trained amino-acid- and descriptor-based MaxEnt models on representative IgH and TCRβ CDR3 repertoires (Fig. 2) and asked which type of model better described test sets of CDR3 sequences set aside from each repertoire, using a nearest- and next-nearest-neighbor amino-acid model as the benchmark (Methods) (Mora et al., 2010). We compared this benchmark to two descriptor models: one that fit similar positional information but with fewer parameters, and one that fit more positional information with a more similar number of parameters. To score these comparisons, we calculated the (logarithm of the) relative probability that each sequence *σ* in the relevant test set belonged to its repertoire according to each of the two models (*M_d_*, descriptor model; *M_a_*, amino-acid model):

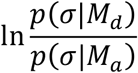

and calculated the percent of sequences for which each model was a better fit. A score of 100% for a given model meant that that model gave a higher probability for every sequence in the test set. As validation, we confirmed that IgH models scored >99% of IgH sequences better than TCRβ models (Fig. 3a, left), and TCRβ models scored >99% of TCRβ sequences better than IgH models (Fig. 3a, right).

**Figure 2.**
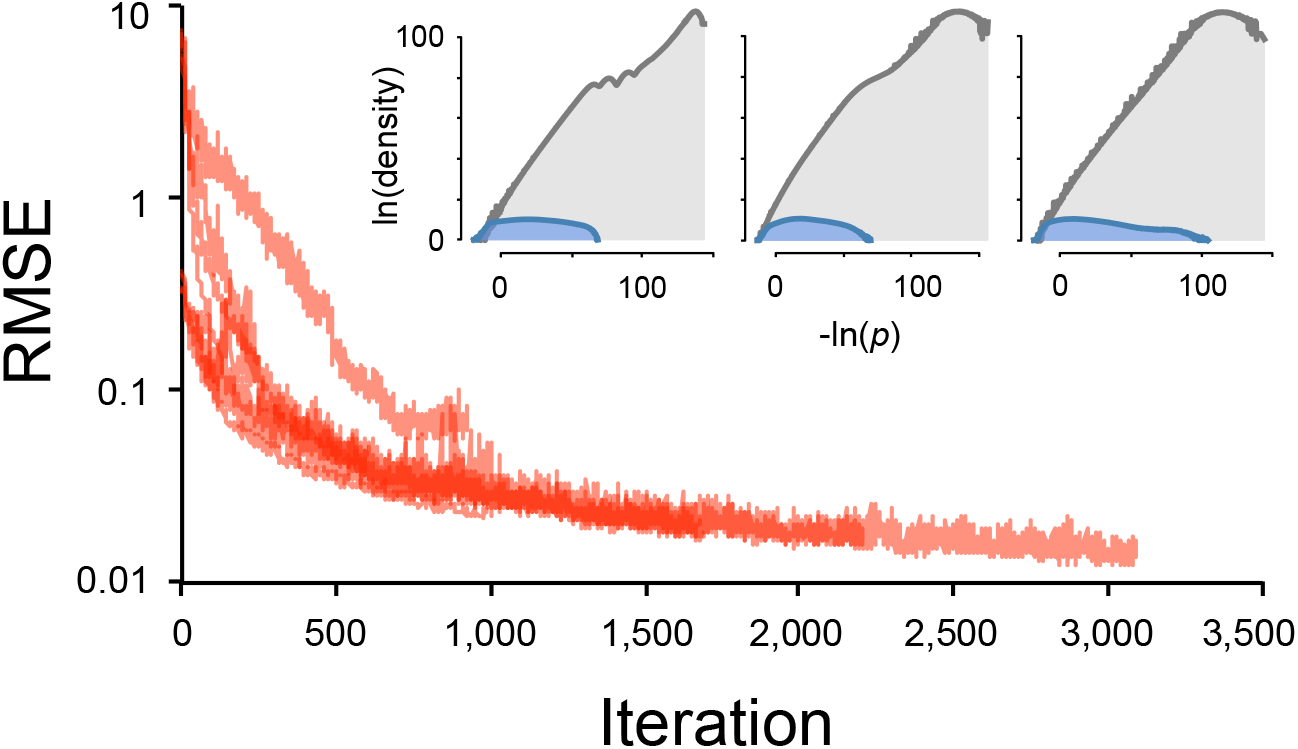
Training and normalization. Distance (root-mean-squared error, RMSE) between training data and model sample as a function of iterations of model training. Data shown is for all models in Tests 1 and 2. Insets, bridge sampling for representative fits showing overlap between model-(blue) and randomly sampled sequences (gray).

**Figure 3.**
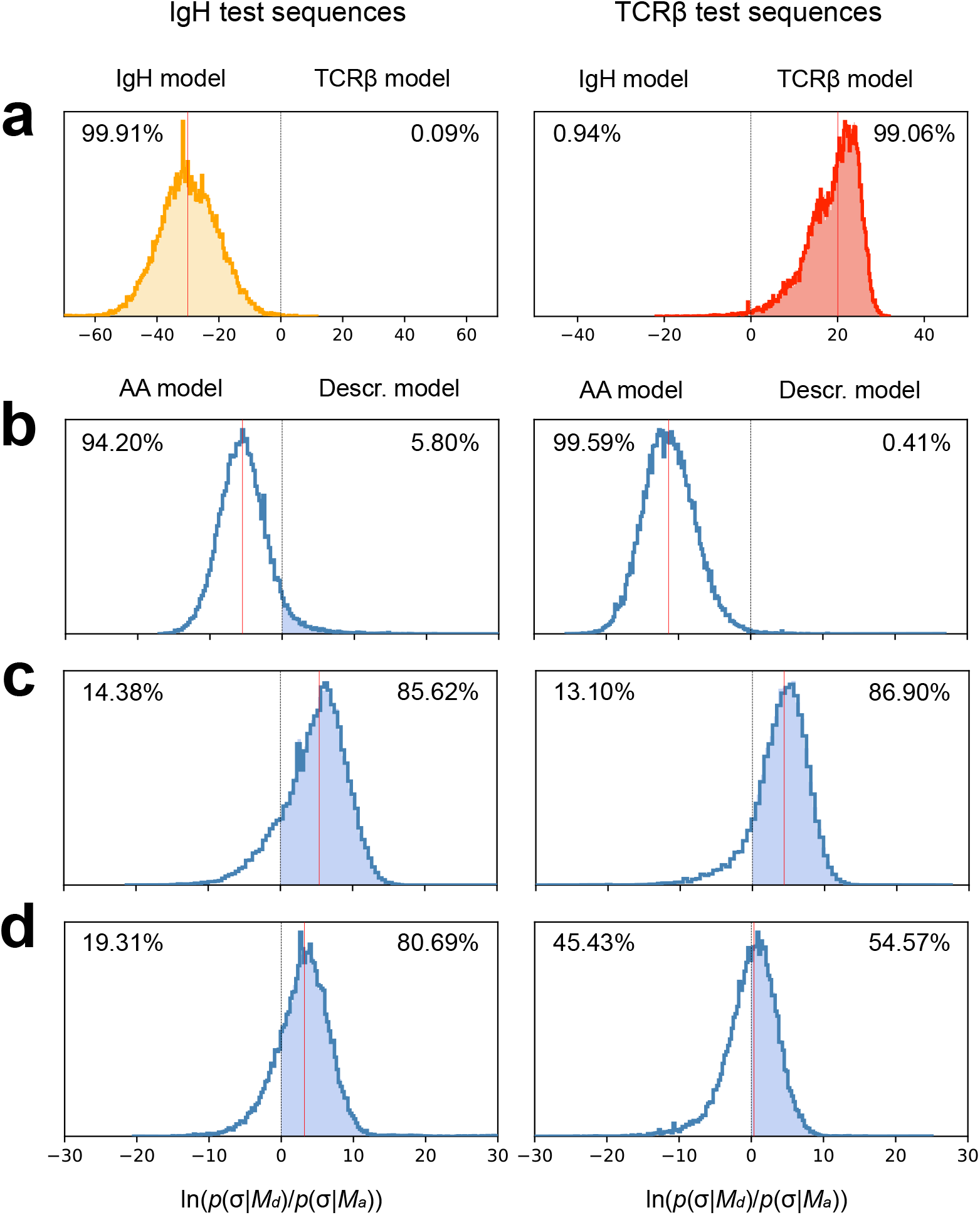
Comparison of amino-acid vs. descriptor models. Head-to-head tests on IgH (left) and TCRβ (right) repertoires; the better performer is shaded green. **(a)** Validation comparison of models of IgH vs. TCRβ repertoires; IgH models strongly prefer IgH sequences (yellow) and TCRβ models strongly prefer TCRβ sequences (red; results shown are for the 325-parameter descriptor models). (b)-(d) Comparisons of an amino-acid model to a descriptor model, both trained/tested on the same training/test set. Density to the left of the vertical dashed line represents sequences for which the amino-acid model gave the higher probability; density to the right (filled) represents higher probability per the descriptor model. Vertical red lines denote medians of the probability densities. **(b)** Test 1: models fitting similar positional information (single positions plus nearest-and next-nearest neighbors); amino-acid models perform better. **(c)** Test 2: models fitting similar numbers of parameters (420 non-length parameters for the amino-acid model vs. 325 for the descriptor model); descriptor models perform better. **(d)** Test 2, continued: amino-acid benchmark model (820 parameters; nearest- and next-nearest neighbors) vs. the descriptor model in (c); descriptor models perform better.

### Test 1: Similar positional information

We first compared models that incorporated similar positional information: single-amino-acid positions and nearest- and next-nearest neighbor pairs (see Methods). The amino-acid models required 2×20^2^=800 parameters to capture the pairwise information vs. just 2×5^2^=50 parameters for the descriptor models (for each of IgH and TCRβ). We predicted that amino-acid models would outperform descriptor models on this test, since for every pair of positions the amino-acid model should have a slight edge, given that descriptors capture only 92% of the variance in amino acids’ biophysical properties. Thus this test was expected to provide an estimate of the cost of swapping alphabets. As predicted, amino-acid models outperformed descriptor models, by a wide margin: 94.2% to 5.8% for IgH (Fig. 3b, left) and 99.6% to 0.4% for TCRβ (Fig. 3b, right). The median sequence had a probability that was ~240 (IgH) and ~87,000 (TCRβ) times as high according to the amino-acid model as according to the descriptor model. For the amino-acid models, sequences from the final samples often contained the canonical CDR3 stems (see Discussion), but these were rare for final samples from these simple descriptor models.!

### Test 2: Similar numbers of parameters

A primary motivation for developing descriptor models is their ability to capture information at a given set of positions with fewer parameters than amino-acid models; the corollary is that for a given number of parameters, descriptor models can capture more positional information. Specifically, for the 400 parameters amino-acid models require to capture information about nearest-neighbor pairs, descriptor models can also capture information about next-nearest-neighbor pairs and cross-loop (Buck, 1992; Weitzner et al., 2015) pairs, both for the stem (or “torso;” see Methods) and the entire CDR3, as well as about consecutive three-amino-acid motifs (*n*=325 non-length parameters for descriptor models vs. 420 for amino-acid models, including the 20 single-amino-acid biases). We therefore first compared 420-parameter amino acid models against 325-parameter descriptor models that fit this additional information.

We expected the descriptor models to outperform these amino-acid models, which, unlike our benchmark amino-acid models, did not fit next-nearest-neighbor pairs, reflecting the utility of additional positional constraints for defining CDR3s. We found that descriptor models outperformed amino-acid models handily, with scores of 85.6% to 14.4% for IgH (Fig. 3c, left) and 86.9% to 13.1% for TCRβ (Fig. 3c, right). The median sequence in the test set was 217- and 82-fold more likely to have been produced by the descriptor model for IgH and TCRβ, respectively. More remarkably, descriptor models also outperformed our benchmark amino-acid models, even though the descriptor models had less than half the parameters (820 vs. 325 non-length parameters), by almost the same margin for IgH, 80.7% to 19.3% (Fig. 3d, left), but by much less for TCRβ, at 54.6% to 45.4% (Fig. 3d, right), leaving the main advantages in this case being coverage and training time. In contrast to the amino-acid models, the familiar start and end motifs (see Discussion) had already been learned in just a few iterations/minutes, requiring just a few hundred sample sequences on which to learn.

### Test 3: Classification

Finally, we sought to test the utility of descriptor models in distinguishing between states of health. As proof of principle, we fit descriptor models on 31 before-and-after IgG+ repertoires (including three replicates) from 14 healthy human volunteers who were administered a seasonal trivalent influenza vaccine (Vollmers et al., 2013). We had previously shown that vaccination leads to prominent changes in both repertoires’ raw and functional diversity (Arora et al., 2018), but withheld diversity measurements from the present study in order to test the discriminatory power of the models in the absence of that additional information. Using stratified 3-fold cross-validation, we found that descriptor models distinguished between pre- and post-vaccination pairs with median AUC of 0.97 (*p*=4×10^-3^; Fig. 4). It is worth noting that applying PCA to the models to reduce them to two dimensions, failed to distinguish between day 0 and day 7, consistent with a lack of necessity for directions of greatest variance to correlate with differences in states of health.

**Figure 4.**
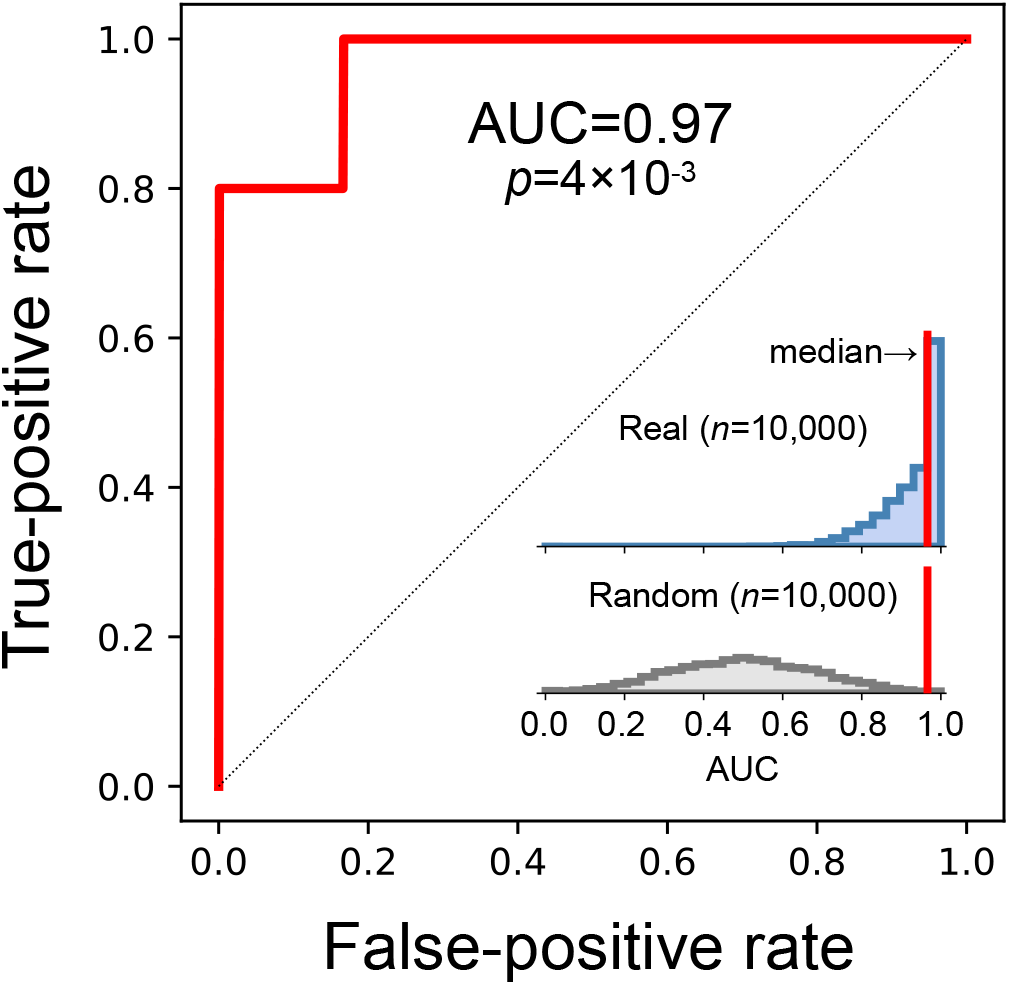
Classification of pre-vs. post-flu vaccination in human subjects. Shown is the median AUC (red) for 10,000 training-test splits using stratified 3-fold cross-validation of an SVM on 31 pre- and post-vaccination samples from the same subjects. Insets show the distributions of AUCs from all 10,000 splits of the real data (blue) and from 10,000 splits in which the data was randomly relabeled, to measure the probability that the median performance could have been the result of chance (gray). Red, median. The p-value is the area in the random-relabeling distribution to the right of the median.

## Discussion

MaxEnt is a powerful method for modeling highly complex systems such as IgH and TCRβ repertoires but exhibits practical limitations related to speed and dimensionality when fit on amino acids using only the standard 20-letter alphabet. Here we demonstrate significant advantages by fitting on biophysical descriptors. We show that appropriate descriptor models can capture more of the information in the repertoire with fewer parameters, and that they can successfully classify health-based states with high accuracy, using the IgG+ B-cell response to influenza vaccination as proof of principle.

A key finding was that descriptor models outperformed amino acid models only once additional positional information was included; when fit on similar positional information—single/overall frequencies and nearest- and next-nearest neighbors—amino-acid models performed better. This finding raises the question of what the relative contributions are of the additional types of positional information fit by the winning descriptor models. There were three additional types of positional information beyond nearest-neighbors: parameters for the stem, cross-pairs, and triples. We chose to fit the stem explicitly because the first and last few amino acids in CDR3s of both IgH and TCRβ are stereotypical, almost canonically beginning with a cysteine (excluded in some definitions), followed by a hydroxylic or small aliphatic amino acid (most often glycine, alanine, or threonine) at the second position, and a basic amino acid (arginine/lysine) at the third position and ending with a methionine or phenylalanine, followed by an aspartate, then valine or tyrosine, and finally tryptophan for IgH, and starting with cysteine, alanine, and a pair of hydrox-ylic or basic amino acids and often ending with glutamate, a variable amino acid, and two aromatic amino acids for TCRβ. These amino acids are important in establishing the stem-loop (or “torso-head”) configuration of CDR3s (Buck, 1992; Weitzner et al., 2015). In IgH and TCRβ the stem is most often encoded by the end of the V gene segment and start of the J, not by the highly variable D gene segment and adjacent non-templated nucleotides (Lefranc et al., 1999); fitting the stem may be allowing the remaining parameters to better fit the more variable region. Fitting cross pairs—the product of descriptor values at the first and last amino acids, second and second-to-last, etc.—was similarly inspired by CDR3s’ stem-loop architecture and may be having a similar benefit. Amino-acid triples are important parts of binding motifs and have been shown to have discriminatory power in IgH in model systems (Sun et al., 2017); it is reasonable that the biophysical patterns they represent also add resolving/discriminatory power. A systematic dissection of these contributions is left for future work.

A further finding was the disparity between the performance of the top descriptor model for IgH, relative to the benchmark amino-acid model, vs. that for TCRβ: the descriptor model scored 80.7% for IgH vs. only 54.6% for TCRβ. A value over 50% indicates that the descriptor model is capturing more information than the amino-acid model, but in the case of TCRβ, the benefit was modest. We considered three possible explanations. First, it is possible that both models captured substantially all of the information present in the training set; however, had this been the case, the models’ final samples would likely have been nearly identical to the training set, and they were not. Second, the additional information in the TCRβ repertoire may not be well captured by the additional positional relationships fit by these models (stem, cross-pairs, triples), but may reside instead in some other relationship(s). Third, the modest benefit may mean that there are isolated (i.e. discontinuous) probability densities in this training set, which the Markov chain used to generate samples (Fig. 1c) has difficulty navigating (van Ravenzwaaij et al., 2018). If so, it may be that somatic hypermutation in the IgH CDR3s bridges probability densities in IgH repertoires that in TCRβ repertoires, which lack somatic hypermutation, remain separate. Conversely, the greater improvement noted for IgH may reflect descriptor models’ ability to detect biophysical similarities among these related sequences, which may be less prominent in TCRβ repertoires but simultaneously difficult to capture in amino-acid models.

The success of descriptor models in correctly discriminating between pre- and post-influenza vaccination suggests potential medical applications. We note that vaccination, like many immunological perturbations, results in systems-as well as sequence-level changes; for example, changes in immunological/repertoire diversity (Jiang et al., 2013; Vollmers et al., 2013). We pre viously showed that the combination of raw and functional diversity, measured with various frequency weightings, can discriminate between pre- and post-vaccination sample pairs with high accuracy, likely in part by detecting clonal expansion with selection (Arora et al., 2018). However, changes in diversity, while potentially useful as part of a screening test, are not sufficiently specific to serve as a general diagnostic modality. The present study shows that even without the powerful discriminatory information that diversity adds, descriptor models are capable of highly sensitive and specific diagnostic discrimination, with high AUC and low p-value from small numbers of subjects and samples. The relatively small number of parameters and these parameters’ relatively straightforward interpretability (compared to, for example, parameters in deep-learning models) suggest that leveraging the statistical biophysics of repertoires’ amino-acid composition is a promising direction for dissecting immune responses for diagnostic and therapeutic purposes. This method is extensible to more or all of IgH or TCRβ, to the complementary chain (IgL/TCRα), and indeed to other proteins or biopolymers, leveraging the power of functional relationships to shrink alphabets while increasing their information density.

## Acknowledgements

This work used the resources allocated via Jetstream cloud service (allocation ID: TG-BI0170094) of Extreme Science and Engineering Discovery Environment (XSEDE), which is supported by National Science Foundation grant number ACI-1548562. Research Computing Group at the High Performance Computing Cluster at Harvard Medical School. The authors would like to thank Dr. Mohammed Al-Quraishi for helpful discussion.

## Methods

### Descriptors

Twenty-six biophysical measurements were previously made on a set of 87 amino acids, which included the standard 20 (Sandberg et al., 1998). We filtered out non-standard amino acids and applied PCA to the standard 20 amino acids (using Python; sklearn.decomposition.PCA library). The top five PCs, which together explained 92% of the observed variance, were each normalized to a mean of 0 and maximum range [-1, 1] and used as biophysical descriptors.

### Data

IgG (Vollmers et al., 2013), memory IgH (DeWitt et al., 2016), and TCRβ (Emerson et al., 2017) CDR3 repertoires were obtained and processed as previously described (Arora et al., 2018). For each dataset in Tests 1 and 2, 500,000 sequences were chosen at random and split 90:10 into training and test sets; for Test 3 all sequences were used.

### Models

MaxEnt models were trained on features’ expectation values, with one parameter per feature (the bias). For Tests 1 and 2, amino-acid models were trained on the observed frequencies of the single amino acids (*n*=20 parameters) and nearest- and next-nearest-neighbor amino-acid pairs (*n*=20^2^×2=800) and the frequencies of CDR3 lengths (*n*=38 for IgH and 26 for TCRβ), following previous reports (Mora et al., 2010). Descriptor models were trained on the frequencies of the single amino acids (*n*=20 parameters), the product of each pair of descriptors at different positions (*n*=5^2^=25 per set), lengths, and, as indicated, on amino-acid frequencies for the first- and last-four amino acids (roughly corresponding to the CDR3 stem or “torso” (North et al., 2011; Finn et al., 2016); *n*=20), the product of each (non-redundant) pair of descriptors at the same position (*n*=(5×4)/2=10), the product of each pair of descriptors for the stem (*n*=25 per set), and the product of descriptors at each three (*n*=5^3^=125). For Test 3, models were trained on the expectation values of each descriptor for the stem (*n*=5) and full-length CDR3 (*n*=5), pairs of descriptors at the same position, cross-loop pairs, and nearest- and next-nearest-neighbor pairs for the full-length CDR3 and the stem (*n*=25 per set), anchoring sequences with an initial cysteine and terminal tryptophan for speed.

Fitting was performed using Metropolis-Hastings Markov-chain Monte Carlo sampling with the acceptance criterion

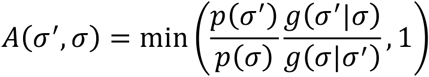

where *σ* is the original sequence and *σ*′ is proposed according to the proposal distribution *g*(*σ*|*σ*′), updating biases via gradient descent using an adaptive step size, using an adaptive burn-in period and autocorrelation time, and a time limit of 24 hours/fit as a stopping condition. Each model was trained for 24 hours on 44 parallel CPUs using the National Science Foundation’s high-performance supercomputing cluster, XSEDE (Towns et al., 2014). To avoid overfitting, we prohibited sample size from exceeding the size of the training set.

### Probabilities

The probability of a sequence *σ* according to a MaxEnt model *M* was calculated as

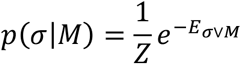

where *E_σ|M_* is the energy of *σ* and the normalization constant *Z* = Σ_*σ*_ *e*^-*E*_*σ|M*_^ was estimated via bridge sampling (Meng and Wong, 1996; Gelman and Meng, 1998) using Harvard Medical School’s high-performance computing cluster.

### Classification (Test 3)

We fit descriptor models on each of the day 0/day 7 before-and-after IgG^+^ repertoire pairs (*n*=31: *n*=17 from day 0, including replicates, and *n*=14 day 7) from the influenza vaccination dataset (Vollmers et al., 2013; Arora et al., 2018) and used a support-vector-machine (SVM) on the final models for classification (excluding length biases, which interact with the normalization constant), using the median area under the receiver-operator-characteristic curve (AUC/ROC) as the quality measure (taken over *n*=10,000 repeats; mean preferred over median given the observed highly skew AUC distributions expected from strong performance with outliers; Fig. 4 top inset), with stratified *k*-fold cross-validation (without oversampling; 17 vs. 14 was considered sufficiently balanced, but see null-model comparison below) to avoid overfitting (for *k*=2, 3, 5, and 10 to confirm robustness) and comparison to the AUC of randomly relabeled data as a null model (also *n*=10,000 repeats) to assess statistical significance. Mann-Whitney U p-value was calculated to test that the two AUC distributions were different. The significance of the AUC was understood as the probability that it could arise from a random classifier by chance; the p-value for significance of the AUC was therefore calculated as the fraction of the area under the null-model distribution to the right of the AUC. Histograms were plotted. All analyses were performed using Python’s numpy and scipy libraries.

## References

Arnaout, R., Lee, W., Cahill, P., Honan, T., Sparrow, T., Weiand, M., Nusbaum, C., Rajewsky, K., and Koralov, S.B. (2011). High-Resolution Description of Antibody Heavy-Chain Repertoires in Humans. PLOS ONE 6, e22365.

Arora, R., Burke, H.M., and Arnaout, R. (2018). Immunological Diversity with Similarity. BioRxiv 483131.

Bialek, W., and Ranganathan, R. (2007). Rediscovering the power of pairwise interactions. ArXiv:0712.4397 [q-Bio].

Buck, C.A. (1992). Immunoglobulin superfamily: structure, function and relationship to other receptor molecules. Semin. Cell Biol. 3, 179–188.

DeWitt, W.S., Lindau, P., Snyder, T.M., Sherwood, A.M., Vignali, M., Carlson, C.S., Greenberg, P.D., Duerkopp, N., Emerson, R.O., and Robins, H.S. (2016). A Public Database of Memory and Naive B-Cell Receptor Sequences. PLoS ONE 11, e0160853.

Dosztányi, Z., and Torda, A.E. (2001). Amino acid similarity matrices based on force fields. Bioinformatics 17, 686–699.

Emerson, R.O., DeWitt, W.S., Vignali, M., Gravley, J., Hu, J.K., Osborne, E.J., Desmarais, C., Klinger, M., Carlson, C.S., Hansen, J.A., et al. (2017). Immunosequencing identifies signatures of cytomegalovirus exposure history and HLA-mediated effects on the T cell repertoire. Nat. Genet. 49, 659–665.

Finn, J.A., Koehler Leman, J., Willis, J.R., Cisneros, A., Crowe, J.E., and Meiler, J. (2016). Im-proving Loop Modeling of the Antibody Complementarity-Determining Region 3 Using Knowledge-Based Restraints. PLoS ONE 11, e0154811.

Gelman, A., and Meng, X.-L. (1998). Simulating Normalizing Constants: From Importance Sampling to Bridge Sampling to Path Sampling. Statistical Science 13, 163–185.

Greiff, V., Weber, C.R., Palme, J., Bodenhofer, U., Miho, E., Menzel, U., and Reddy, S.T. (2017). Learning the High-Dimensional Immunogenomic Features That Predict Public and Private Anti-body Repertoires. J. Immunol. 199, 2985–2997.

Hellberg, S., Sjöström, M., Skagerberg, B., and Wold, S. (1987). Peptide quantitative structure-activity relationships, a multivariate approach. J. Med. Chem. 30, 1126–1135.

Jiang, N., He, J., Weinstein, J.A., Penland, L., Sasaki, S., He, X.-S., Dekker, C.L., Zheng, N.-Y., Huang, M., Sullivan, M., et al. (2013). Lineage structure of the human antibody repertoire in response to influenza vaccination. Sci Transl Med 5, 171ra19.

Kaplinsky, J., Li, A., Sun, A., Coffre, M., Koralov, S.B., and Arnaout, R. (2014). Antibody repertoire deep sequencing reveals antigen-independent selection in maturing B cells. Proc. Natl. Acad. Sci. U.S.A. 111, E2622–2629.

Kim, S., Thiessen, P.A., Bolton, E.E., Chen, J., Fu, G., Gindulyte, A., Han, L., He, J., He, S., Shoemaker, B.A., et al. (2016). PubChem Substance and Compound databases. Nucleic Acids Res. 44, D1202–1213.

Lefranc, M.P., Giudicelli, V., Ginestoux, C., Bodmer, J., Müller, W., Bontrop, R., Lemaitre, M., Malik, A., Barbié, V., and Chaume, D. (1999). IMGT, the international ImMunoGeneTics database. Nucleic Acids Res. 27, 209–212.

Marks, D.S., Colwell, L.J., Sheridan, R., Hopf, T.A., Pagnani, A., Zecchina, R., and Sander, C. (2011). Protein 3D Structure Computed from Evolutionary Sequence Variation. PLOS ONE 6, e28766.

Meng, X.-L., and Wong, W.H. (1996). Simulating ratios of normalizing constants via a simple identity: A theoretical exploration. Statistica Sinica 6, 831–860.

Mora, T., Walczak, A.M., Bialek, W., and Callan, C.G. (2010). Maximum entropy models for antibody diversity. PNAS 107, 5405–5410.

North, B., Lehmann, A., and Dunbrack, R.L. (2011). A new clustering of antibody CDR loop con-formations. J. Mol. Biol. 406, 228–256.

Ostmeyer, J., Christley, S., Rounds, W.H., Toby, I., Greenberg, B.M., Monson, N.L., and Cowell, L.G. (2017). Statistical classifiers for diagnosing disease from immune repertoires: a case study using multiple sclerosis. BMC Bioinformatics 18, 401.

Ostmeyer, J., Christley, S., Toby, I.T., and Cowell, L.G. (2019). Biophysicochemical motifs in T cell receptor sequences distinguish repertoires from tumor-infiltrating lymphocytes and adjacent healthy tissue. Cancer Res canres.2292.2018.

Parameswaran, P., Liu, Y., Roskin, K.M., Jackson, K.K.L., Dixit, V.P., Lee, J.-Y., Artiles, K.L., Zompi, S., Vargas, M.J., Simen, B.B., et al. (2013). Convergent antibody signatures in human dengue. Cell Host Microbe 13, 691–700.

van Ravenzwaaij, D., Cassey, P., and Brown, S.D. (2018). A simple introduction to Markov Chain Monte–Carlo sampling. Psychon Bull Rev 25, 143–154.

Robins, H.S., Srivastava, S.K., Campregher, P.V., Turtle, C.J., Andriesen, J., Riddell, S.R., Carlson, C.S., and Warren, E.H. (2010). Overlap and Effective Size of the Human CD8+ T Cell Receptor Repertoire. Science Translational Medicine 2, 47ra64–47ra64.

Russ, W.P., Lowery, D.M., Mishra, P., Yaffe, M.B., and Ranganathan, R. (2005). Natural-like function in artificial WW domains. Nature 437, 579–583.

Sandberg, M., Eriksson, L., Jonsson, J., Sjöström, M., and Wold, S. (1998). New chemical descriptors relevant for the design of biologically active peptides. A multivariate characterization of 87 amino acids. J. Med. Chem. 41, 2481–2491.

Seno, F., Trovato, A., Banavar, J.R., and Maritan, A. (2008). Maximum entropy approach for deducing amino Acid interactions in proteins. Phys. Rev. Lett. 100, 078102.

Stephenson, J.D., and Freeland, S.J. (2013). Unearthing the root of amino acid similarity. J. Mol. Evol. 77, 159–169.

Sun, Y., Best, K., Cinelli, M., Heather, J.M., Reich-Zeliger, S., Shifrut, E., Friedman, N., Shawe-Taylor, J., and Chain, B. (2017). Specificity, Privacy, and Degeneracy in the CD4 T Cell Receptor Repertoire Following Immunization. Front. Immunol. 8.

Susko, E., and Roger, A.J. (2007). On reduced amino acid alphabets for phylogenetic inference. Mol. Biol. Evol. 24, 2139–2150.

Towns, J., Cockerill, T., Dahan, M., Foster, I., Gaither, K., Grimshaw, A., Hazlewood, V., Lathrop, S., Lifka, D., Peterson, G.D., et al. (2014). XSEDE: Accelerating Scientific Discovery. Computing in Science & Engineering 16, 62–74.

Vollmers, C., Sit, R.V., Weinstein, J.A., Dekker, C.L., and Quake, S.R. (2013). Genetic measurement of memory B-cell recall using antibody repertoire sequencing. Proc. Natl. Acad. Sci. U.S.A. 110, 13463–13468.

Walter, K.U., Vamvaca, K., and Hilvert, D. (2005). An active enzyme constructed from a 9-amino acid alphabet. J. Biol. Chem. 280, 37742–37746.

Weitzner, B.D., Dunbrack, R.L., and Gray, J.J. (2015). The origin of CDR H3 structural diversity. Structure 23, 302–311.

Yeo, G., and Burge, C.B. (2004). Maximum entropy modeling of short sequence motifs with applications to RNA splicing signals. J. Comput. Biol. 11, 377–394.

